# The effect of facial expression on contrast sensitivity: a behavioural investigation and extension of Hedger, Garner, & Adams (2015)

**DOI:** 10.1101/432229

**Authors:** Abigail L. M. Webb, Paul B. Hibbard

## Abstract

It has been argued that rapid visual processing for fearful face expressions is driven by the fact that effective contrast is higher in these faces compared to other expressions, when the contrast sensitivity function is taken into account (Hedger, Garner, & Adams, 2015). This proposal has been upheld by data from image analyses, but is yet to be tested at the behavioural level. The present study conducts a traditional contrast sensitivity task for face images of various facial expressions. Findings show that visual contrast thresholds do not differ for different facial expressions We re-conduct analysis of faces’ effective contrast, using the procedure developed by Hedger, Garner, & Adams (2015), and show that higher effective contrast in fearful face expressions relies on face images first being normalised for RMS contrast. When not normalised for RMS contrast, effective contrast in fear expressions is no different, or sometimes even lower, compared to other expressions. These findings are discussed in relation to the implications of contrast normalisation on the salience of face expressions in behavioural and neurophysiological experiments, and also the extent that natural physical differences between facial stimuli are masked during stimulus standardisation and normalisation.

## Introduction

Fearful facial expressions are particularly salient to the human visual system, receiving preferential allocation of attentional resources, and inhibiting this attention from relocating to different stimuli^[1–4]^. This attentional effect is also found when fearful faces appear in peripheral vision^[5–6]^. When fearful expressions compete with salient noise stimuli for visual awareness, with the face and noise presented to different eyes, they break suppression faster compared to neutral faces^[7–8]^, and are associated with increased activity in subcortical threat-processing regions even when observers report not having observed a face^[5–6, 9]^. These findings converge on the notion that the human visual system has evolved specific visual neural mechanisms that enable rapid identification of fearful expressions. This concept is reminiscent of LeDoux’s^[10]^ ‘quick and dirty pathway’ for processing environmental information necessary for successful threat-avoidance.

A visual stimulus might be selectively processed for two reasons: because it is semantically and meaningfully relevant, or because its configuration is somehow congruent with low-level mechanisms in early vision that allow for it to be rapidly and efficiently processed. In terms of the threat bias for fearful faces, this means that fearful faces may be prioritised because of their emotional relevance, or their low-level image properties. The latter, low-level approach has been a particular focus within visual psychophysics, where studies have shown that it is specifically the low spatial frequency information in fearful faces that gives rise to the saliency effects associated with fearful expressions^[1, 11–12]^.. Low frequency components of fear expressions are thought to undergo rapid processing via low-frequency-sensitive subcortical pathways that directly access the amygdala^[11–12]^. Such findings are interpreted as evidence of visual mechanisms that selectively respond to signals present in fearful faces^[13–14]^. Hedger, Garner, & Adams^[15]^ propose an equally low-level, but directionally different account, arguing that stimulus properties characteristic of fearful expressions ensure strong responses in the early stages of visual processing^[7, 15]^. Gray^[7]^ and Hedger^[15]^ make use of this sensory bias hypothesis to explain how perceptual biases for fear expressions may be accounted for by the way in which their physical attributes are well matched to the sensitivity of early visual processing, as opposed to the recruitment of attentional mechanisms that preferentially respond to expressions of fear^[15, 16]^. Here, a distinction is made between stimulus detection that arises from attentional processes, and that which occurs pre-attentively. Here, Hedger and colleagues^[15]^ implicate the contrast sensitivity function in the threat bias.

Hedger, Garner and Adams^[15]^ compared the Fourier amplitude spectra of images of fearful and neutral faces, since both the overall contrast and the spatial frequency content of images are known to modulate stimulus salience, with the human visual system being most efficient at detecting information around 3–5 cpd. They assessed the effective contrast of images by multiplying the Fourier amplitude spectra of face stimuli by a standard measure of the contrast sensitivity function, based on the Modelfest data set^[17]^. This approach quantifies effective contrast as the product of the image amplitude, and the visual system’s sensitivity, at each spatial frequency. They found that fearful faces, when matched for RMS contrast, were higher in effective contrast, and therefore better matched for the contrast sensitivity function compared to neutral faces. This finding was accounted for by the higher degree of contrast energy in midrange spatial frequencies, where the contrast sensitivity function peaks, in fearful faces. The effect was found to be consistent across several commonly used face databases including the Karolinska Directed Emotional Faces^[18]^, Radbound Faces^[19]^, Ekman and Friesen^[20]^, Montreal set of facial displays (MSFDE)^[21]^ and NimStim databases^[22]^. Data from their image analyses support the notion that biases for fearful expressions are driven at least in part by their sensory efficacy.

However, there remain several elements of this approach that are not addressed by Hedger, Garner and Adams^[15]^. The first relates to the behavioural evidence in support of the efficacy-account. Evidence for a role of preconscious processing of threat-information is provided by the results of experiments using the continuous flash suppression (CFS) paradigm, but does not directly measure expression-related effects on contrast sensitivity. That is, while the more rapid detection of fearful faces in CFS experiments is consistent with their greater effective contrast, we might also expect differences in contrast sensitivity to different facial expressions to be evident more directly. Specifically, if it is the case that the Fourier amplitude spectrum of fearful faces is well matched to the human contrast sensitivity function, we should expect to observe an increase in contrast sensitivity, reflected by decreased contrast thresholds, for fearful faces. The second issue is that prior to the transformation of face images, Hedger and colleagues^[15]^ normalised face images for their luminance and RMS contrast, such that they were identical on these measures at the physical level. While this is a commonly employed technique in psychophysical studies, performed to reduce contrast-driven differences in stimulus salience, the process of attributing the aggregate physical contrast to all facial stimuli may mask naturally occurring differences in contrast between expressions in a way that could obscure results. Normalising images of natural scenes ensures a degree of consistency between images’ physical and perceived salience. However, the same may not be true when applied to face images. O’Hare & Hibbard^[23]^ show inconsistencies between images’ physical and apparent contrast when there are differences in amplitude spectrum, when these stimuli are matched for RMS content. Given the uncertain effects of normalisation on the physical and perceived salience of facial stimuli, it is reasonable to question the degree to which normalisation influences results from both image analyses and behavioural paradigms. In particular, any consistent differences in RMS contrast across facial expressions would be expected either to increase, or cancel out, differences in sensitivity that can be attributed to differences in effective contrast.

To address these questions, we conducted a replication of the image analyses performed by Hedger, Garner and Adams^[15]^. We included face stimuli that are physically matched for RMS contrast, but also faces that were physically unmatched, such that they contain natural differences in both physical and apparent contrast. Furthermore, we conducted a traditional contrast sensitivity task in order to psychophysically test predictions from Hedger’s image analysis. Here, we employed facial expressions as opposed to sinusoidal grating stimuli to measure expression-related differences in contrast sensitivity. An important feature of this latter study is that it directly addresses the association between face expression and contrast sensitivity at the behavioural level.

## Materials and Methods

### Participants

Eighteen (15 female, 3 male) participants took part in the study. All participants were informed of the nature of the study and provided written informed consent prior to the study beginning. The University of Essex Ethics Committee approved the employed experimental procedures. All participated in the experiment as part of a credited research module assessment, or in exchange for monetary reward. All participants had normal to corrected vision.

### Stimuli and Apparatus

Stimuli were grayscale images of 16 individuals, 8 male and 8 females, taken from the Karolinska Directed Emotional Faces set^[18]^. Face images included internal features only, and included 4 emotional expressions of neutral, fear, anger and happiness. Though Hedger and colleagues^[15]^ refer to only fearful and neutral expressions, we included an additional two expressions so as to include positively and negatively-valenced comparisons. All individual faces were presented in their normal, upright form, and in a phase scrambled format. Phase scrambled versions of the face images were used as a control measure, providing versions of faces whose configural content was disrupted but low level statistical properties preserved. An example is shown in Figure 1. Phase scrambling was performed using MATLAB fast Fourier transform functions. Contrast thresholds were determined using an adaptive staircase technique (see under Procedure, below). Stimuli were presented using a VIEWPIXX 3D monitor (52cm X 29cm), viewed from a distance of 65 cm. The stimulus size of faces was 5.5 degrees. The screen resolution was 1920×1080 pixels, with a refresh rate of 120Hz and an average luminance of 50 cdm^-2^. Each pixel subtended 1.43 arc min. Stimuli were presented at 10-bit resolution. Participants’ responses were recorded using the RESPONSEPixx response box. Stimuli were generated and presented using MATLAB and the Psychophysics Tool box extensions^[24–26]^.

**Figure 1:**
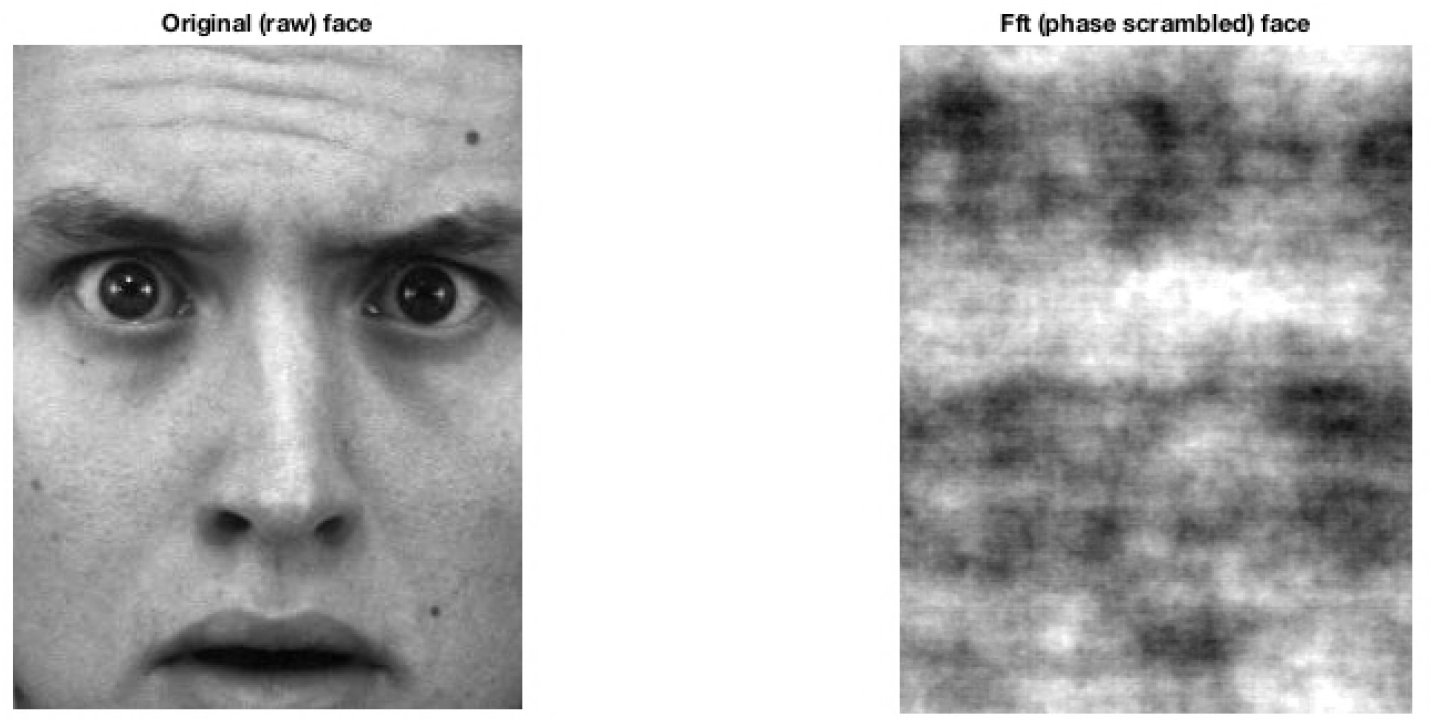
Image of a fearful face expression extracted from the KDEF database (left), and a fast Fourier transform (FFT) of the same image (right). The same face image is therefore shown in its raw, normal format (left), and in a phase-scrambled format (right). Both images share the same amplitude spectrum (contrast-spatial frequency association), but the random phase structure assigned to the right image distributes this information differently.

### Procedure

Participants were tested individually in a quiet room and informed prior to the experiment that the study was concerned with face perception.

As a 2AFC location task, participants’ objective was to indicate, using 1 of 2 buttons on a RESPONSEPixx response box whether the target face image appeared to the left or right of centre. The beginning of each trial commenced with the face stimulus on the left or right side of the screen. Participant responses determined the onset of the next trial. The proportion of times that the participant correctly indicated the location of the stimulus was recorded for all face stimuli.

The adaptive staircase method was used to establish the Michelson contrast required for correct detection (75% of the time) for each expression stimulus. Here, the starting contrast level for each expressions’ staircase began at 0.01 Michelson contrast. According to the up-down rule^[27]^, Michelson contrast was increased by one initial step of 0.005 proceeding 1 incorrect observer response, thus boosting stimulus visibility. Conversely, 3 correct observer responses triggered a decrease in Michelson contrast, initially by 0.005. The overall staircase length was 70 trials, where the initial step size (0.005 Michelson) halved after 17, 35 and 52 trials. 4 experimental blocks were completed, and the 280 trials for each combination of expression and phase scrambling were combined to create a single psychometric function.

## Results

The proportions of participants’ correct responses for each expression, at each contrast level, were used to create a psychometric function. A cumulative Gaussian function was fit to this data using the Palemedes toolbox^[28]^ and used to determine a contrast detection threshold for each expression in its normal and manipulated (scrambled) formats. This 75% contrast detection threshold was defined as the contrast required for the participant to correctly identify the location of the face stimulus on 75% of trials. These results are plotted in Figure 2.

**Figure 2:**
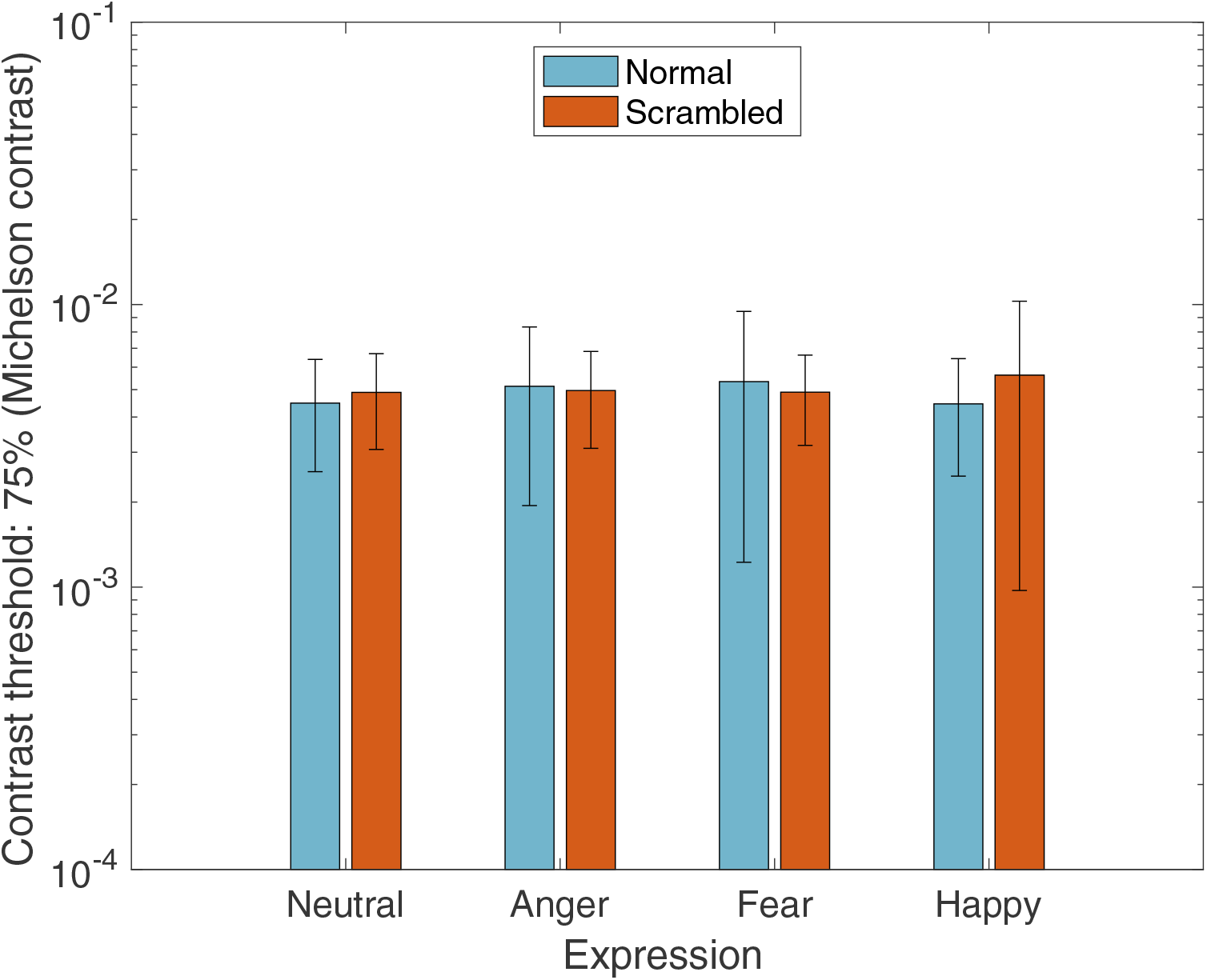
Visual contrast thresholds for neutral, fearful, angry and happy facial expressions. Error bars show standard error values. Faces are unfiltered. Fearful face expressions are not associated with lower visual contrast thresholds, contrary to what might be predicted from Hedger et al., (2015). Error bars represent ±1 standard error of the mean.

A 4 Emotion (neutral, anger, fear, happy) x 2 Manipulation (normal, scrambled) within subjects ANOVA revealed no significant effects of expression (*F*(3, 51) =. 26, *p*=.85, ηp ^2^=.01), or manipulation (*F*(1, 17)= .13, *p*= .72, ηp ^2^ =.01), and no significant expression x manipulation interaction (*F*(3, 51) = 1.20, *p =*. 32, ηp ^2^ =. 06). Analyses were repeated for contrast thresholds that were calculated using the RMS contrast of face stimuli. A 4 Emotion (neutral, anger, fear, happy) x 2 Manipulation (normal, scrambled) within subjects ANOVA revealed no significant effect of expression (*F*(3, 51) =. 42, *p* =. 73, ηp^2^ =. 02), or manipulation (*F*(1, 17) =. 07, *p =*. 79 ηp ^2^ =. 004), and no significant expression x manipulation interaction (*F*(3, 51) = 1.23, *p* =. 31, ηp ^2^ =. 07). These findings show that visual contrast thresholds do not vary between face expressions, nor are these findings different according to the two contrast metrics used here (Michelson and RMS). The absence of an expression-related effect on contrast sensitivity provides evidence against Hedger and colleagues’ ^[15]^ original claim that fear expressions (compared to neutral faces) exploit the contrast sensitivity function. In an attempt to understand the inconsistency between the present behavioural data, and that generated from image analyses, we conducted the same measure of faces’ effective contrast as that performed by Hedger, Garner and Adams ^[15]^ and extended this to include expressions of anger, happiness and disgust, including a condition where all face images had been either normalised for RMS contrast (as was the procedure for Hedger and colleagues^[15]^) or non-normalised, such that face images were analysed in their raw format, containing possible natural variations in physical contrast.

### Image analyses

Hedger and colleagues^[15]^ calculated the effective contrast for face images extracted from 5 face databases: Nimstim, KDEF, Radbound, Montreal and Ekman and Friesen face sets. Stimuli were cropped to include internal features only and normalised for RMS contrast prior to analyses.

Here, we calculate effective contrast for the 16 KDEF face images used in our experimental study, referring to the same procedure described by Hedger, Garner and Adams^[15]^. First, Fourier amplitude spectra were calculated for each face image. From the ModelFest dataset^[17]^, we extracted visual contrast thresholds for 10 stimulus parameters. These corresponded to Gabor stimuli, ranging from 1.12–30 cycles/degree. A smooth curve was fit to the average threshold (over 4 repetitions and all observers in the ModelFest dataset) using a cubic spline. The resulting contrast sensitive distribution was then multiplied by the Fourier amplitude spectrum for each face image to establish each face’s effective contrast. Figure 3 shows an example of the procedure for calculating effective contrast for the 16 face images used in the present contrast sensitivity study. To extend our analysis, effective contrast was measured for face images across 4 of the face databases employed by Hedger, Garner and Adams^[15]^, with the exception of the Ekman & Friesen face set^[20]^.

**Figure 3:**
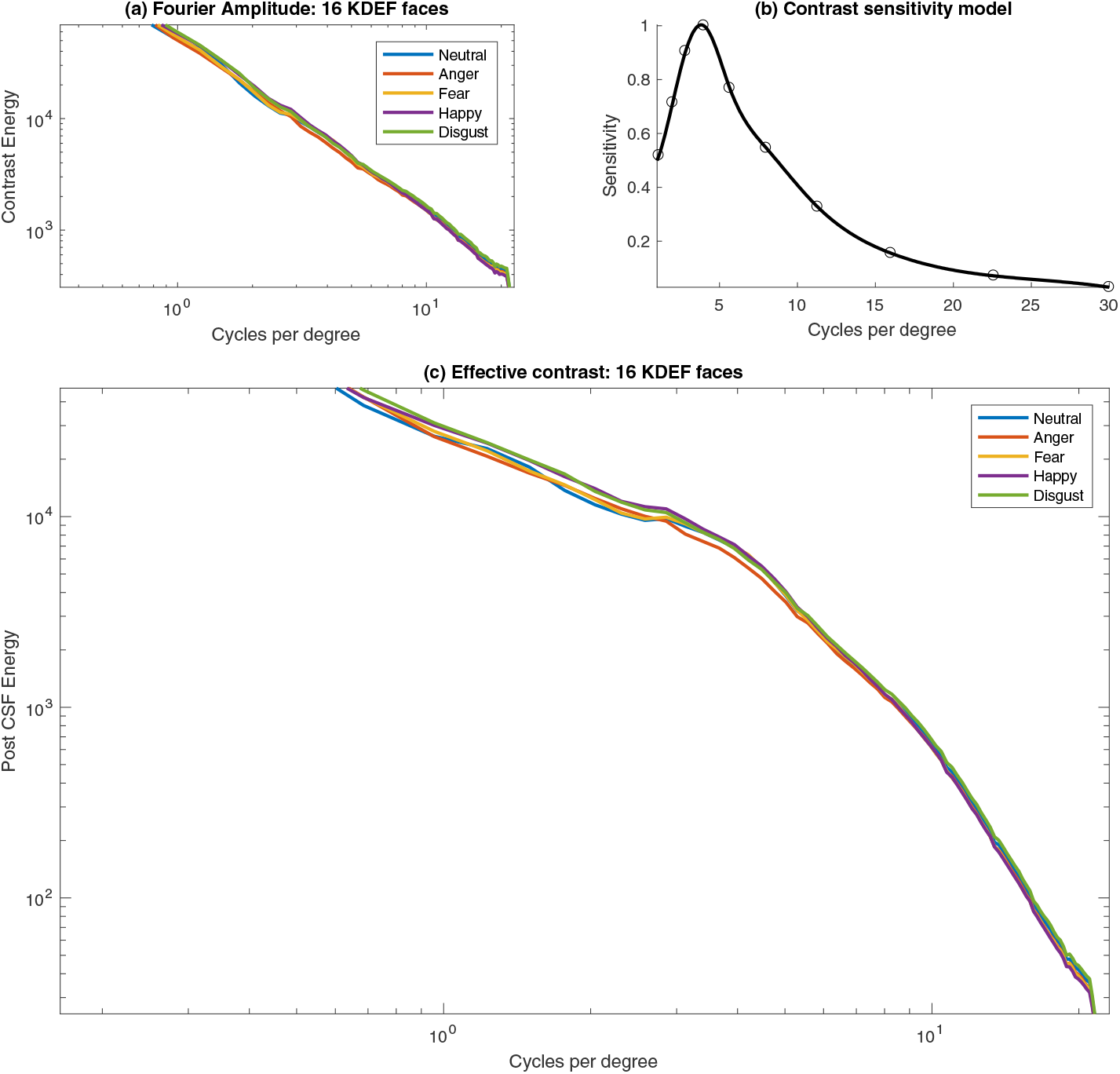
(a) The mean amplitude spectrum for each of the five expressions. *(a)* The contrast sensitivity function based on the ModelFest data. (c) The effective contrast, obtained by multiplying the original amplitude function by the contrast sensitivity function. This method for calculating effective contrast replicates that used by Hedger, Garner and Adams^[15]^.

As outlined by Hedger and colleagues^[15]^ the overall estimate of effective contrast for each face image was obtained by summing contrast across spatial frequency after application of the contrast sensitivity model. All face images were analysed in two conditions: after they had been normalised for RMS contrast (according to Hedger and colleagues^[15]^), and also in their raw form, such that no contrast normalisation had taken place. In the RMS-matched analysis, the RMS contrast of each face was set to be equal to that of the image with the *lowest* contrast in each set. It is for this reason that the RMS-matched stimuli have an overall lower effective contrast. All face images depict forward-facing actors displaying one of 5 expressions (neutral, anger, fear, happy or disgust), cropped to include internal features only. The average effective contrast for each facial expression, compared across the 5 face image samples, including the experimental stimuli for the present contrast sensitivity study, is displayed in Figure 4.

**Figure 4:**
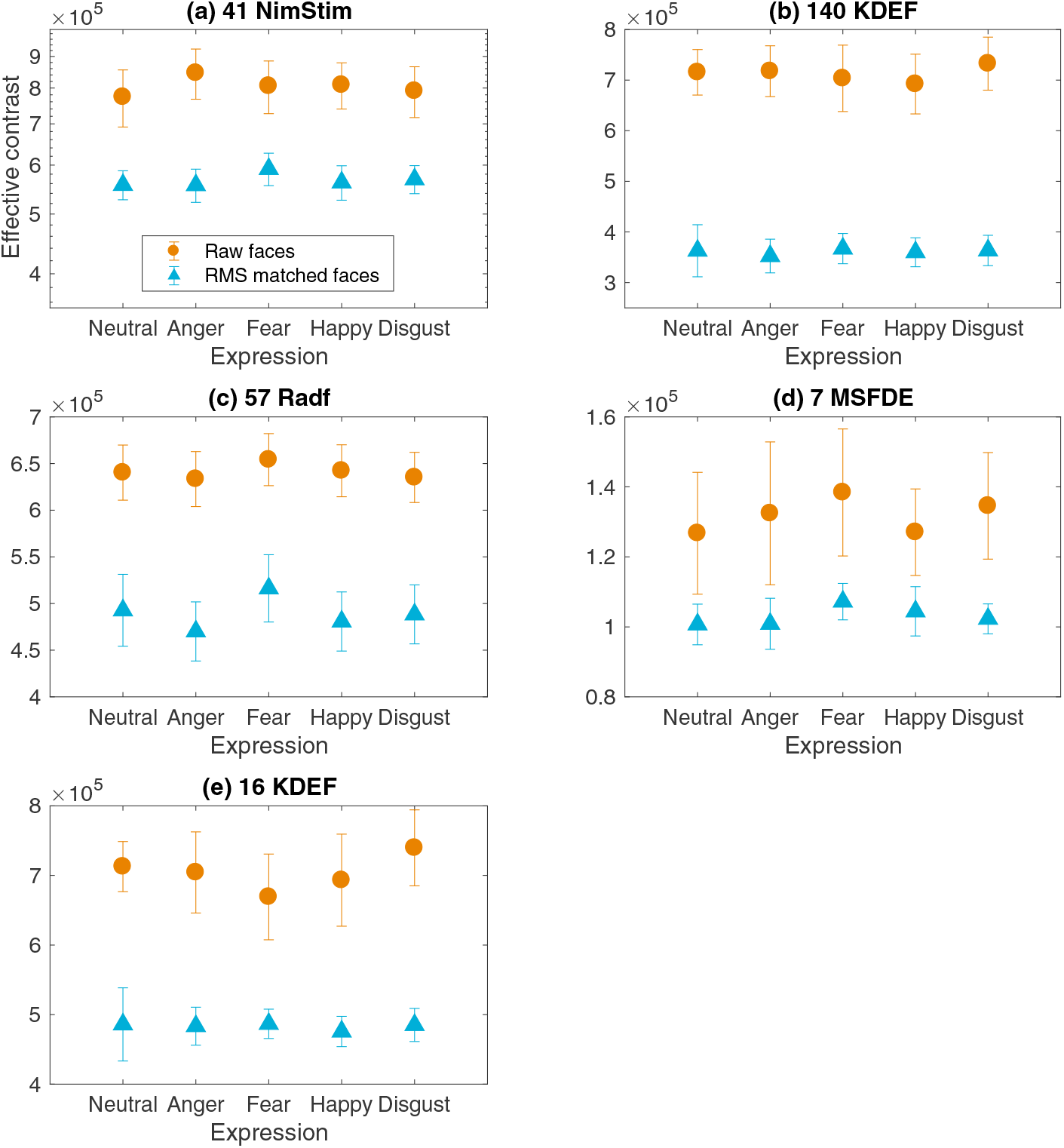
Effective contrast for neutral faces, and anger, fear, happiness and disgust expressions, measured for raw faces (circle data) and the same faces normalised for RMS contrast (triangle data). Effective contrast measures were performed across 4 samples of face images, including the NimStim (a), KDEF (b), Radbound (c), MSFDE (d), face sets employed by Hedger et al., (2015), and for the 16 KDEF faces used in the present contrast sensitivity study (e). Error bars represent ±1 standard error of the mean.

#### 42 NimStim face images

Effective contrast for neutral, angry, fearful, happy and disgust NimStim faces are shown in Figure 4 (a), and summarised in Table 1 (a). Paired comparisons measured the extent that effective contrast associated with fear expressions differed compared to neutral, anger, happy and disgust expression counterparts. When face images were first normalised for RMS contrast, effective contrast belonging to fearful NimStim faces was significantly higher compared to neutral expressions. This finding replicates that observed by Hedger, Garner and Adams^[15]^, whereby RMS normalised fear expressions were found to be significantly higher in effective contrast compared to neutral expressions. Our data extend this finding to show that RMS normalised fear expressions not only contain significantly higher effective contrast than neutral counterparts, but that their effective contrast is also higher than that for angry, happy and disgust expressions. Alternatively, when effective contrast was measured for the same raw NimStim faces that had *not* been normalised for RMS contrast, fear expressions were significantly higher in effective contrast compared to neutral expressions, but significantly lower in effective contrast compared to angry expressions. The effective contrast for 41 raw NimStim fearful faces did not significantly differ from that belonging to happy and disgust expressions.

**Table 1:**
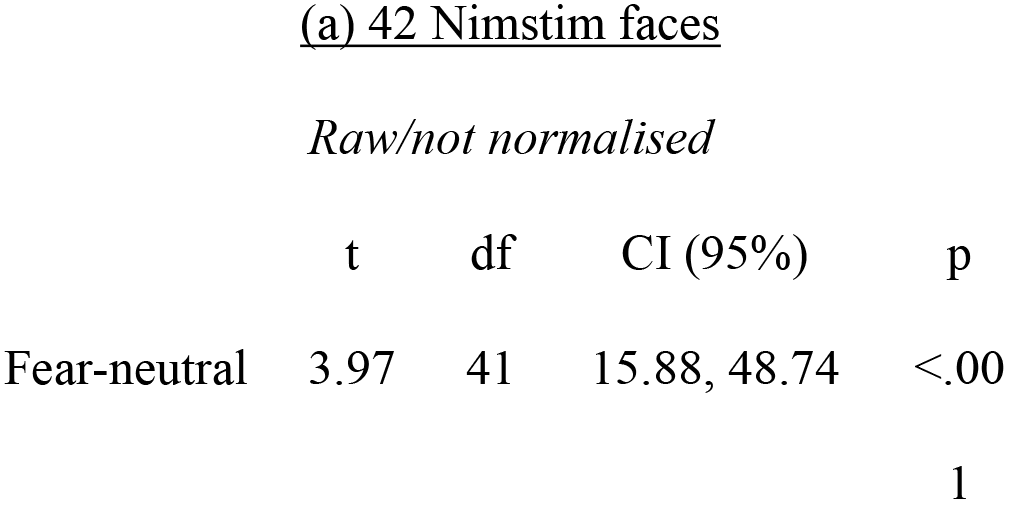
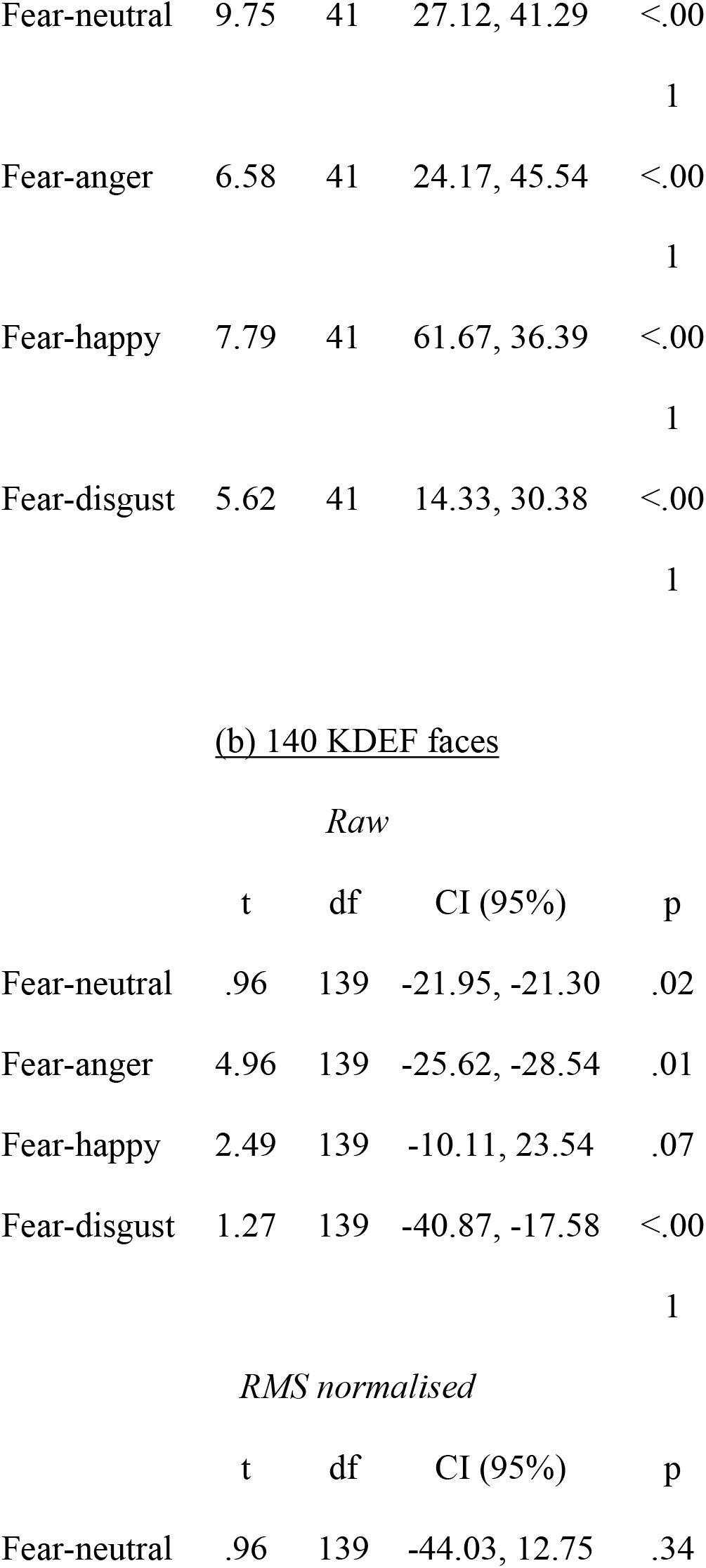
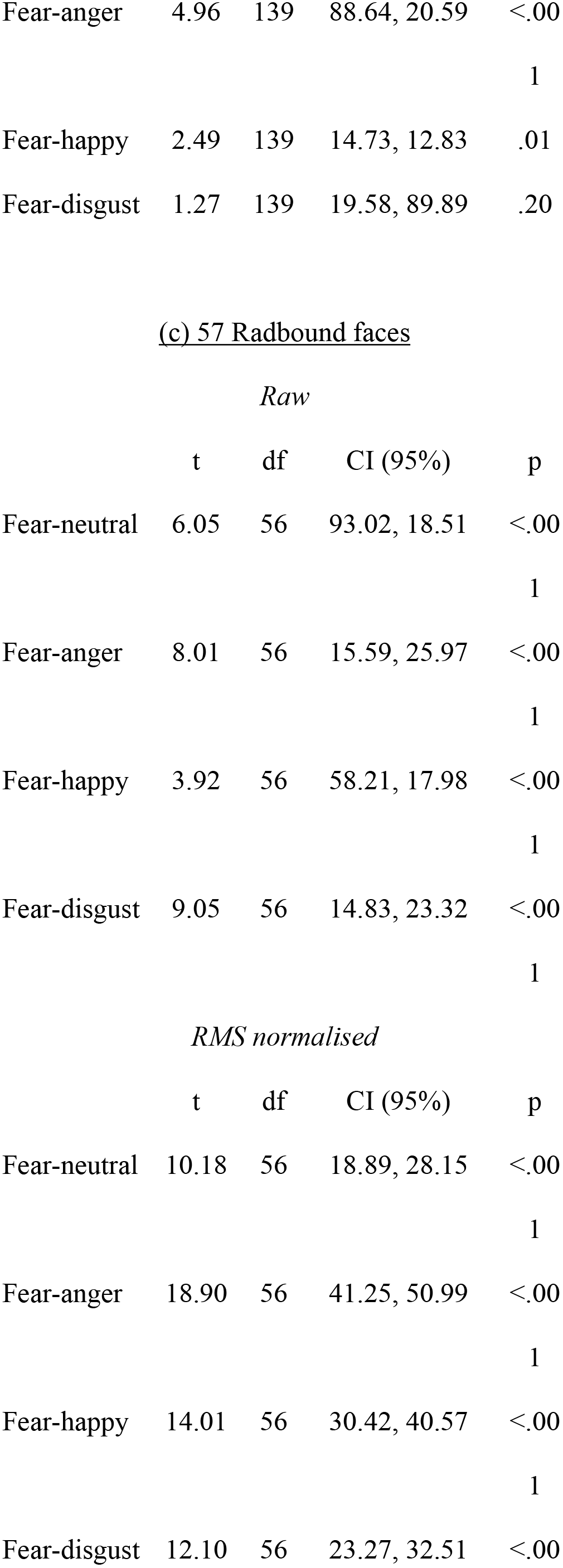
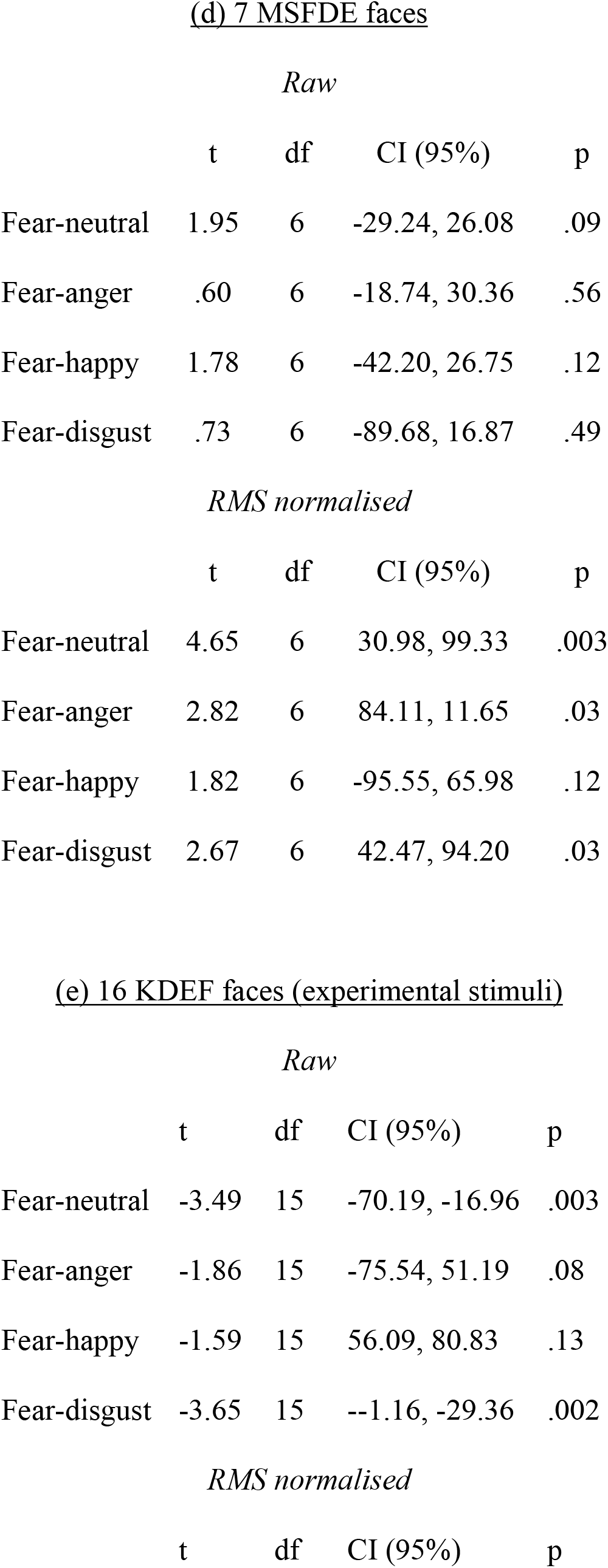
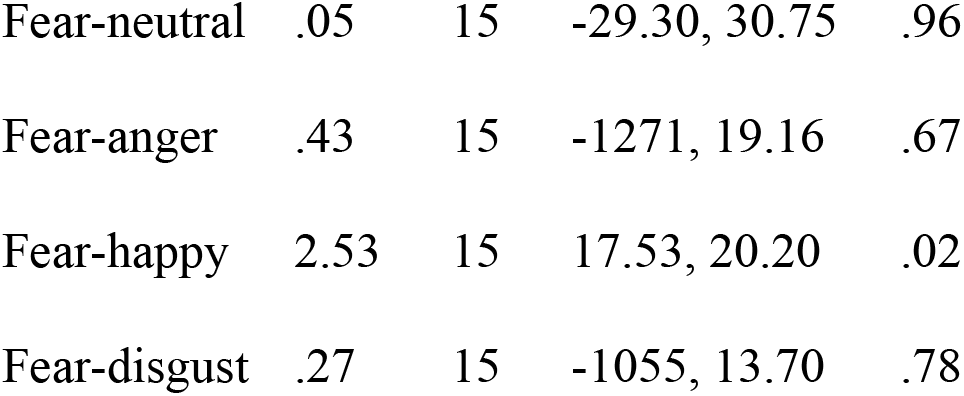
Effective compared between fear and counterpart expressions. Comparisons are made when faces are raw, and thus not normalised for contrast, and also when they are matched for RMS contrast. Measures are performed for 4 databases (a-d), and experimental stimuli used in the present behavioural study (e).

For 42 raw (not normalised) NimStim faces, RMS contrast was calculated across the 5 face expressions in order to understand how natural differences in physical contrast might influence differences between expressions effective contrast according to whether or not they have been normalised for RMS contrast. Fearful Nimstim faces naturally contained significantly less RMS contrast compared to anger and happy expressions, and did not differ compared to neutral and disgust faces. These data are illustrated in Figure 5, and summarised in Table 2 (a).

**Figure 5:**
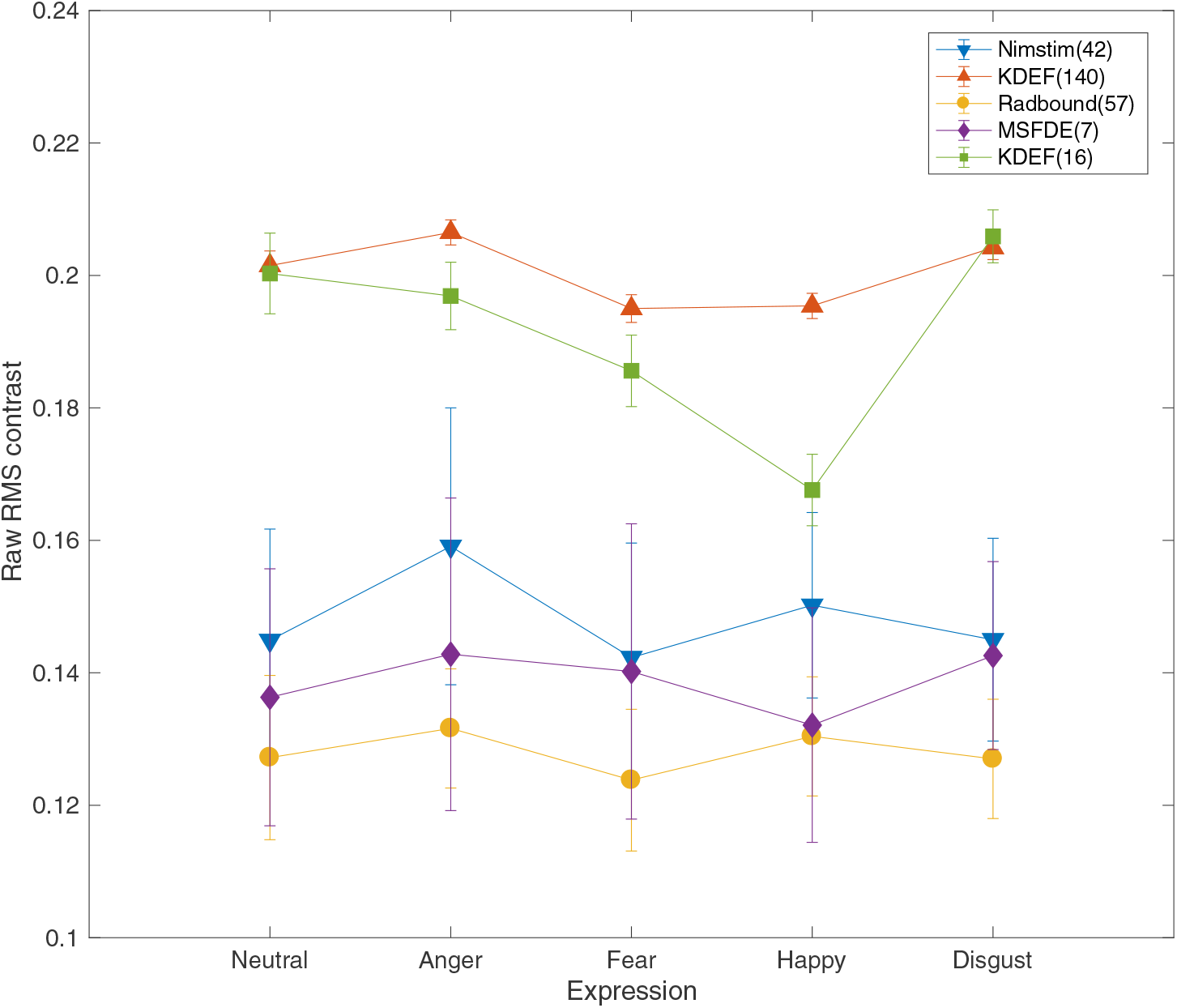
RMS contrast for face expressions *before* faces are subjected to contrast normalisation i.e. when kept in raw format. RMS contrast for 5 expressions is measured across the 5-database face samples used to calculate faces’ effective contrast. Error bars represent ±1 standard error of the mean.

**Table 2:**
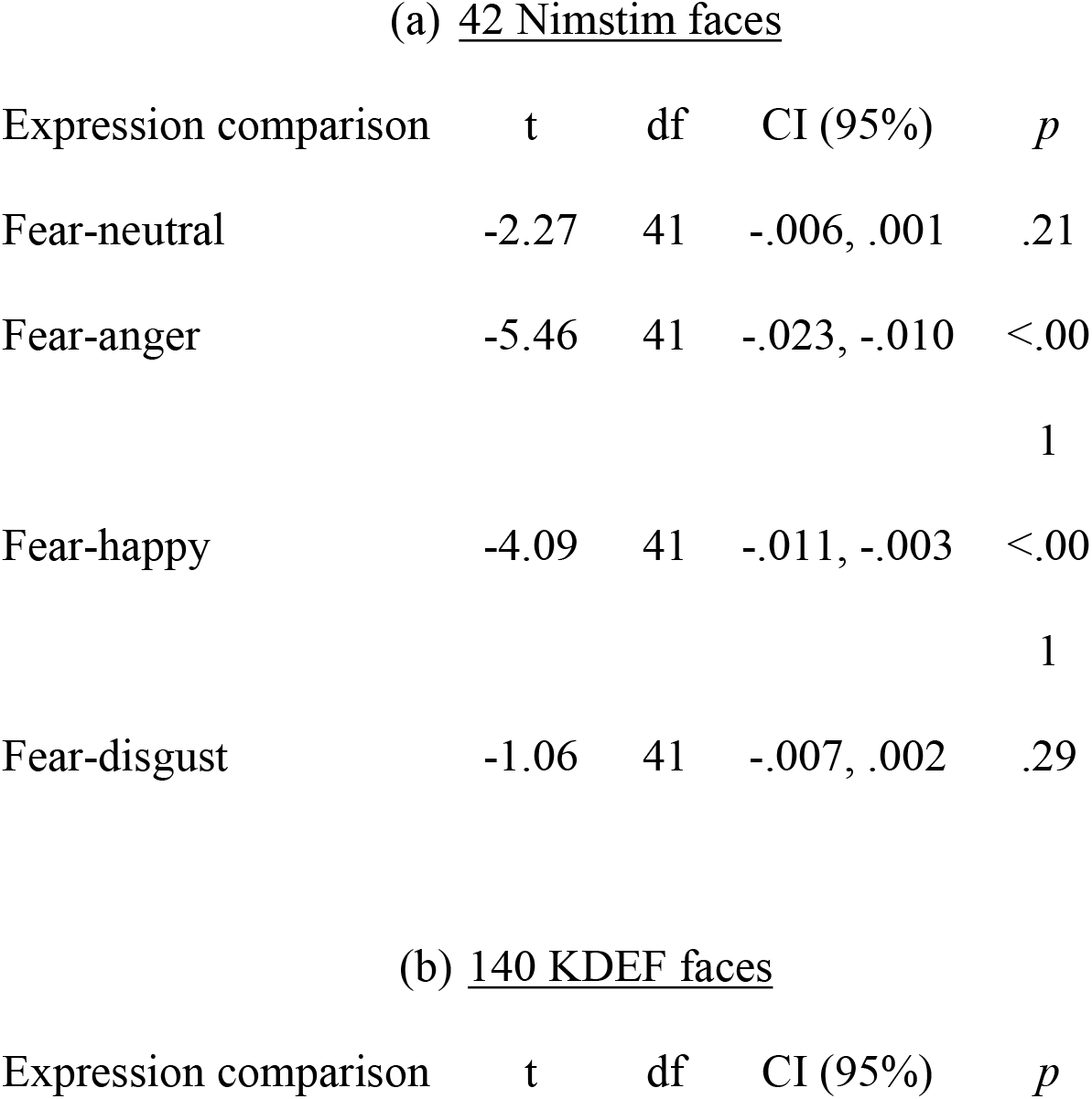
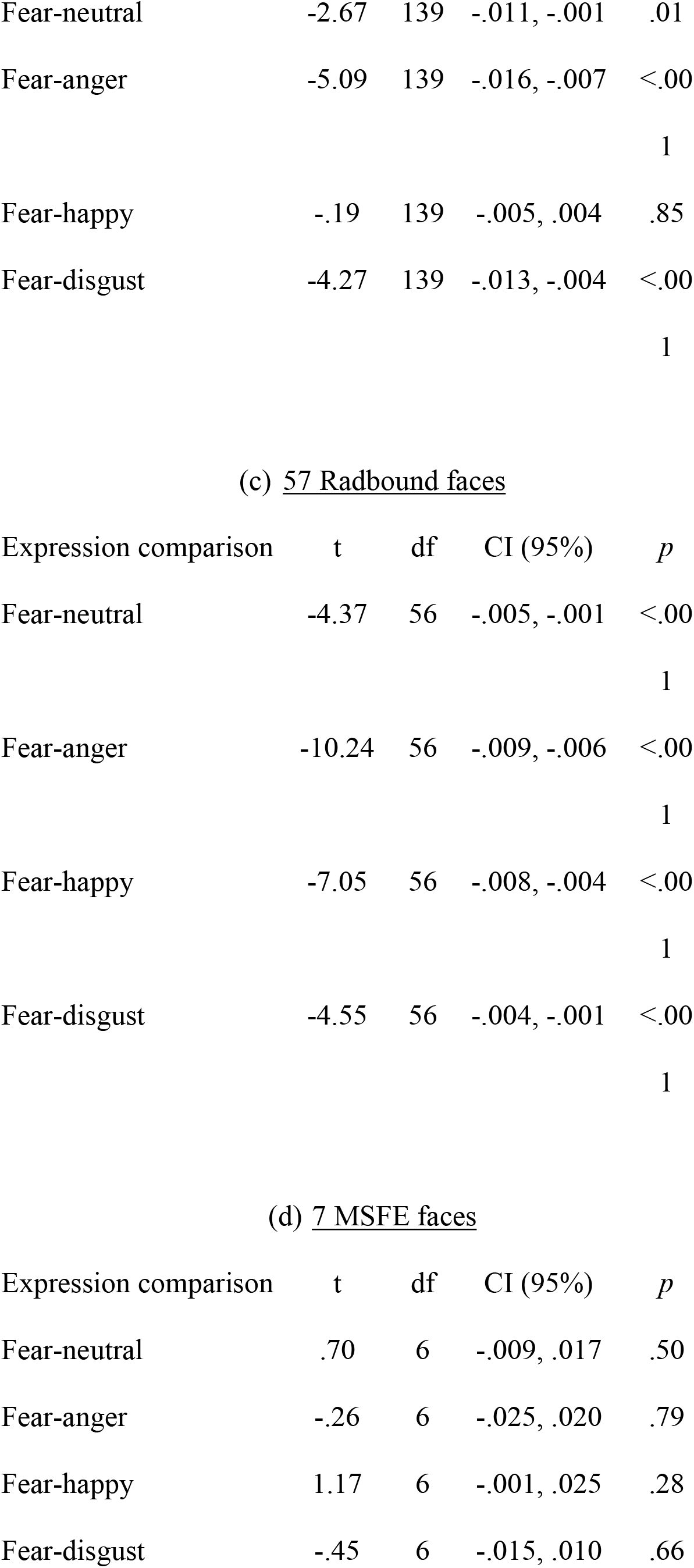
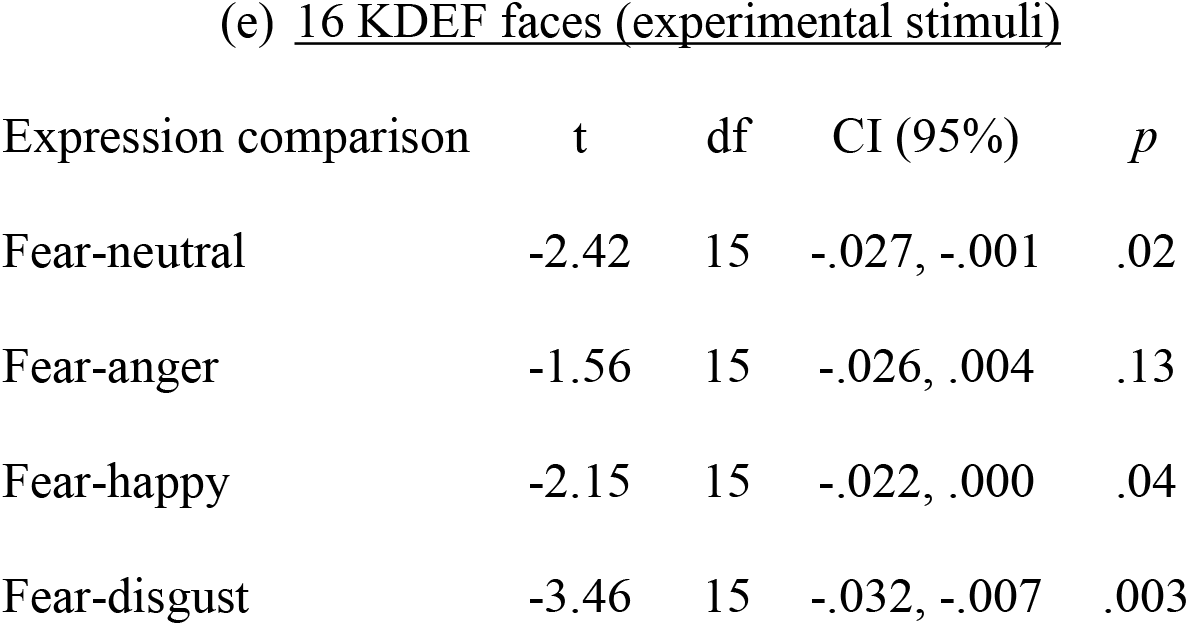
Differences between RMS contrast in raw fear expressions and 4 emotion counterparts (neutral, anger,happy, disgust). Fear comparisons are measured across all 4 databases (a-d), and also for the experimental stimuli used in the present contrast sensitivity study (e).

#### 140 KDEF face images

Effective contrast for neutral, angry, fearful, happy and disgust KDEF faces are shown in Figure 4 (b), and summarised in Table 1 (b). Paired comparisons measured differences in effective contrast between fearful faces and their expression counterparts. For KDEF face images normalised for RMS contrast, fear expressions were significantly higher in effective contrast compared to both angry and happy expressions. Effective contrast did not differ between fear expressions and disgust or neutral faces when normalised for RMS contrast. Alternatively, when effective contrast was measured for the same face images, but in their raw form, fearful KDEF faces were significantly *lower* in effective contrast compared to neutral, angry and disgust expressions, and did not differ compared to happy expressions.

For 140 raw KDEF faces, RMS contrast was calculated across the 5 expressions. Fearful KDEF faces naturally contained significantly less RMS contrast compared to neutral, angry and disgust expressions, and did not differ compared to happy expressions. These data are illustrated in Figure 5, and summarised in Table 2 (b).

#### 57 Radbound face images

Effective contrast for neutral, angry, fearful, happy and disgust Radbound faces are shown in Figure 4 (c), and summarised in Table 1 (c). Paired comparisons measured differences in effective contrast between fearful faces and their expression counterparts. For face images normalised for RMS contrast, fear expressions were significantly higher in effective contrast compared to neutral faces, but also compared to all other expressions. These effects were also true when effective contrast was calculated for the same faces in raw form, thus not normalised for RMS contrast. These findings in particular require further discussion, presented in the following section.

For 57 raw Radbound faces, RMS contrast was calculated across the 5 expressions. Fearful Radbound faces naturally contained significantly less RMS contrast compared to all other face expressions. These data are illustrated in Figure 5, and summarised in Table 2 (c).

#### 7 Montreal (MSFDE) face images

Effective contrast for neutral, angry, fearful, happy and disgust Montreal faces are shown in Figure 4 (d), and summarised in Table 1 (d). Paired comparisons measured differences in effective contrast between fearful faces and their expression counterparts. When face images were normalised for RMS contrast, effective contrast did not differ for fear faces compared to any other face expressions. Alternatively, when effective contrast was measured for the same faces, but in their raw form, fearful Montreal faces were significantly higher in effective contrast compared to neutral, angry and disgust expressions, and did not differ significantly from happy expressions.

For 7 raw Montreal faces, RMS contrast was calculated across the 5 expressions. RMS contrast for fearful Montreal faces did not differ significantly compared to any other face expression. These data are illustrated in Figure 5, and summarised in Table 2 (d).

#### 16 KDEF face images (experimental stimuli)

Effective contrasts for the experimental face stimuli used in our contrast sensitivity study are shown in Figure 4 (e), and summarised in Table 1 (e). Here, effective contrast is measured for experimental face images that were *not* normalised for RMS contrast, and also for versions of the same faces that had been normalised for RMS contrast. Paired comparisons measured differences in effective contrast between fearful faces and their expression counterparts. When faces were normalised for RMS contrast, effective contrast was significantly higher in fear expressions compared to happy expressions, and did not significantly differ compared to neutral, anger or disgust faces. Alternatively, for the same faces that were not normalised for RMS contrast, such that they were analysed in the same format as they were presented to observers, fear expressions were significantly lower in effective contrast compared to both neutral and disgust expressions, and did not differ significantly from angry or happy expressions.

For the 16 raw KDEF faces used in the present contrast sensitivity study, RMS contrast was calculated across the 5 expressions. Fearful faces naturally contained significantly less RMS contrast compared to neutral and disgust faces, more RMS contrast compared to happy faces, and did not differ significantly compared to angry expressions. These data are illustrated in Figure 5, and summarised in Table 2 (e).

Together, data from the present contrast sensitivity study showed that visual contrast thresholds are not influenced by differences between images of facial expressions. Namely, fearful expressions portrayed by face images did not enhance observers’ contrast sensitivity; as was predicted by findings from Hedger, Garner and Adams^[15]^. Fearful expressions, according to image analyses by Hedger, Garner and Adams^[15]^ are higher in effective contrast, and thus well tuned to contrast sensitivity processing. This proposal was driven by data from image analyses measuring differences in effective contrast between fear and neutral face images that had been normalised for RMS contrast. The stimuli used in the present study were raw face images that were not normalised for physical contrast in any way. We replicate measures of effective contrast used by Hedger, Garner and Adams ^[15]^ to establish the extent that CSF advantages exclusive to fear expressions might be driven by the effect of contrast normalisation on the effective contrast of faces. A general trend across the present image analyses is that greater effective contrast in fear expressions appears to rely on face images having been first normalised for RMS contrast. This was the case for both the experimental stimuli used in the present CSF study, and for the face sets used by Hedger and colleagues^[15]^.

## Discussion

A widely accepted view in the threat bias literature is that fearful face expressions possess a special status in the human visual system, due to their low level image content^[1, 3–4, 7, 29]^. Hedger, Garner and Adams^[15]^ recently showed that visibility, or salience, associated with fear expressions is predicted by their effective contrast content; the extent that the Fourier amplitude of fear expressions, compared to neutral faces, exploits the contrast sensitivity function. In the present study, we conducted a traditional contrast sensitivity task to test whether higher effective contrast purported for fear expressions is associated with lower visual contrast thresholds at the behavioural level. Here, we measured contrast sensitivity for facial stimuli of 5 raw face expressions. No expression-related differences were observed across visual thresholds, as was predicted based on data from Hedger, Garner and Adams^[15]^.

Specifically, a decrease in visual thresholds for fearful expressions was not observed. Greater effective contrast unique to fear expressions (when compared to neutral faces) was found only for face images that had been normalised for RMS contrast. In order to investigate whether the use of contrast normalisation in Hedger, Garner and Adams’ ^[15]^ may have driven effective contrast effects that in its absence were not replicated by our contrast sensitivity study, we repeated calculations of effective contrast using the same procedure employed by Hedger, Garner and Adams^[15]^. Here, effective contrast was calculated for images of face expressions both when they were normalised for RMS contrast, as was performed by Hedger, Garner and Adams^[15]^, but also when the same faces had not been normalised for physical contrast. These analyses were performed for the NimStim, KDEF, Montreal (MSFDE) and Radbound face sets used by Hedger and colleagues^[15]^, and also for the 16 KDEF face images used as the experimental stimuli in the present contrast sensitivity study. Importantly, our findings replicate those of Hedger, Garner and Adams^[15]^, showing that fear expressions normalised for RMS contrast are significantly higher in effective contrast than neutral counterparts. We extend this finding to show that this is also true when fearful faces are compared to other face expressions. However, when the same faces were analysed in their raw form (i.e. when they were not normalised for physical contrast), this effect of fear diminishes for Nimstim, KDEF, and Montreal face images. These findings indicate that the process of normalising face stimuli significantly increases the effective contrast in fearful face expressions, where naturally these faces tend not to differ in effective contrast compared to other facial expressions, or indeed possess a lower effective contrast. An important finding to discuss here is the absence of this contrast normalisation effect for face images taken from the Radbound face database. Here, we observed that Radbound fear expressions normalised for contrast were significantly higher in effective contrast compared to neutral faces, as well as other expressions; an effect that is consistent with that observed by Hedger, Garner and Adams^[15]^. However, this effect did not diminish when images were not normalised for contrast, as was observed for all other face samples. Radbound face images were included in the present study on the basis that they were included in the original study by Garner and Adams^[15]^. Details of the image processing used to create and standardise these actor photographs includes white-balance correction^[19]^. This process adjusts raw image data in order to remove certain unrealistic and biased appearances, such as those incurred under different lightning conditions^[30–31]^. It is important to note that database production information for KDEF and NimStim face sets do not refer to any image processing related to white-balance correction, or contrast normalisation^[32, 22]^. No information about image processing is provided for the Montreal image database^[21]^. It may therefore be that contrast and luminance information in ‘raw’ Radbound face images had already been subjected to some degree of normalisation when they were created. In sum, the present study performed a traditional contrast sensitivity task to address the proposal that fearful faces exploit the contrast sensitivity function, and as a result undergoes efficient visual processing^[15]^. Together, these findings suggest that contrast normalisation –a standard procedure in psychophysical studies- significantly influences the physical composition of face stimuli in a way that can be expected to influence their perceived salience under both experimental and neurophysiological conditions.

**Data can be made publicly available via Dryad.**

